# Transposase assisted tagmentation of RNA/DNA hybrid duplexes

**DOI:** 10.1101/2020.01.29.926105

**Authors:** Bo Lu, Liting Dong, Danyang Yi, Chenxu Zhu, Meiling Zhang, Xiaoyu Li, Chengqi Yi

## Abstract

Tn5-mediated transposition of double-strand DNA has been widely utilized in various high-throughput sequencing applications. Here, we report that the Tn5 transposase is also capable of direct tagmentation of RNA/DNA hybrids *in vitro*. As a proof-of-concept application, we utilized this activity to replace the traditional library construction procedure of RNA sequencing, which contains many laborious and time-consuming processes. Results of activity of transposase assisted RNA/DNA hybrids co-tagmentation (termed “ATRAC-seq”) are comparable to traditional RNA-seq methods in terms of gene number, gene body coverage and gene expression analysis; at the meantime, ATRAC-seq enables a one-tube library construction protocol and hence is more rapid (within 8 h) and convenient. We expect this tagmentation activity on RNA/DNA hybrids to have broad potentials on RNA biology and chromatin research.

## Introduction

Transposases exist in both prokaryotes and eukaryotes and catalyze the movement of defined DNA elements (transposon) to another part of the genome in a “cut and paste” mechanism (1–3). Taking advantage of this catalytic activity, transposases are widely used in many biomedical applications: for instance, an engineered, hyperactive Tn5 transposase from *E. coli* has been utilized in an *in vitro* double-stranded DNA (dsDNA) tagmentation reaction to achieve rapid and low-input library construction for next-generation sequencing (4–9). In addition, Tn5 was also used for *in vivo* transposition of native chromatin to profile open chromatin, DNA-binding proteins and nucleosome position (“ATAC-seq”) (10). While Tn5 has been broadly adopted in high-throughput sequencing, bioinformatic analysis and structural studies reveal that it belongs to the retroviral integrase superfamily that act on not only dsDNA but also RNA/DNA hybrids (for instance, RNase H). Despite the distinct substrates, these proteins all share a conserved catalytic RNase H-like domain (see Figure 1a) (11–14). Given their structural and mechanistic similarity, we attempted to ask whether or not Tn5 is able to catalyze tagmentation reactions to RNA/DNA hybrids (see Figure 1b), in addition to its canonical function of dsDNA transposition. In this study, we tested this hypothesis and found that indeed Tn5 possesses *in vitro* tagmentation activity towards both strands of RNA/DNA hybrids. As a proof of concept, we apply such activity of transposase-assisted RNA/DNA hybrids co-tagmentation (ATRAC-seq) to achieve rapid and low-cost RNA sequencing starting from total RNA extracted from 10,000 to 100 cells. We find that ATRAC-seq data are comparable to conventional RNA-seq results in terms of detected gene numbers, gene expression measurement and gene body coverage, at the same time it avoids many laborious and time-consuming steps in traditional RNA-seq experiments. Such Tn5-assisted tagmentation of RNA/DNA hybrids could have broad applications in RNA biology and chromatin research.

**Figure 1.**
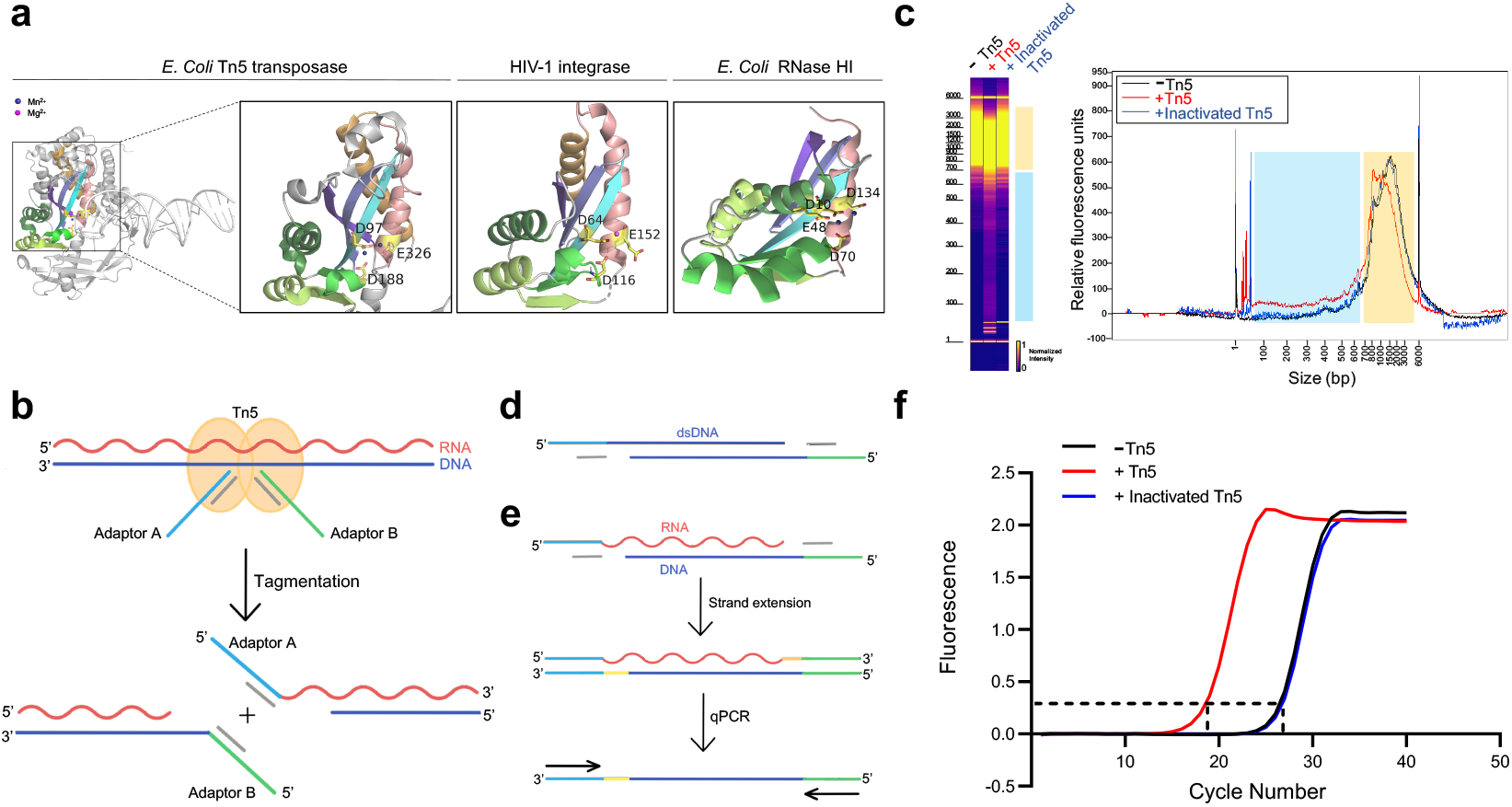
Tn5 transposome has direct tagmentation activity on RNA/DNA hybrid duplexes. **(a)** Crystal structure of a single subunit of *E. coli* Tn5 Transposase (PDB code 1MM8) complexed with ME DNA duplex, and zoom-in views of the conserved catalytic core of Tn5 transposase, HIV-1 integrase (PDB code 1BIU), and *E. coli* RNase HI (PDB code 1G15), all of which are from the retroviral integrase superfamily. Active-site residues are shown as sticks, and the Mn^2+^ and Mg^2+^ ions are shown as deep blue and magenta spheres. **(b)** Schematic of Tn5-assisted tagmentation of RNA/DNA hybrids. **(c)** Gel pictures (left) and peak pictures (right) represent size distributions of RNA/DNA hybrid fragments after incubation without Tn5 transposome, with Tn5 transposome, and with inactivated Tn5 transposome. The blue and orange patches denote small and large fragments, respectively. **(d)** Schematic of the product of *in vitro* tagmentation reaction of the canonical dsDNA substrate. **(e)** Workflow of conversion of tagged RNA/DNA hybrids into amplifiable DNA sequences. **(f)** qPCR amplification curve of tagmentation products of samples with Tn5 treatment, with inactivated Tn5 treatment, or without Tn5 treatment. Average Ct values of two technical replicates are 18.06, 26.25 and 26.41, respectively.

## Results

To test whether Tn5 transposase has tagmentation activity on RNA/DNA hybrids, we prepared RNA/DNA duplexes by performing mRNA reverse transcription. We first validated the efficiency of reverse transcription and the presence of RNA/DNA duplexes using a model mRNA sequence (~1,000 nt) as template (see Figure S1a). We then subjected the prepared RNA/DNA hybrids from 293T mRNA to Tn5 transposome, heat-inactivated Tn5 transposome and a blank control (without Tn5), respectively (see Methods). The hybrids were then recovered and their length distribution was analyzed by Fragment Analyzer (see Figure 1c). Comparing with the heat-inactivated Tn5 sample or the blank control sample, the Tn5 transposome sample exhibited a modest but clear smear signal corresponding to small fragments ranging from ~30-650 base-pair (bp) (the blue patches in Figure 1c). Consistent with the fragmentation event, we also observed a down shift of large fragments ranging from ~700-4000 bp (the orange patches in Figure 1c). In addition, the fragmentation efficiency increased in a dose-dependent manner with the transposome, suggesting that fragmentation of RNA/DNA hybrids is dependent on Tn5 (see Figure S1b).

We next asked whether RNA/DNA hybrids are tagged by Tn5. For a canonical dsDNA substrates, the staggered tagmentation of Tn5 results in a 9 bp gap between the nontransferred strand and the target DNA (see Figure 1d). We anticipate that a similar *in vitro* tagmentation reaction to RNA/DNA hybrids generates a structure with adaptors ligated to the 5’ ends of both RNA and DNA strands and gaps at the 3’ ends (see Figure 1e). If such a structure is present, we would be able to convert it into an amplifiable DNA sequence by reverse transcription from the target DNA into this gap, followed by extension synthesis of the attached adaptor sequence by strand displacement (see Figure 1e). We chose *Bst* 3.0 DNA polymerase, which demonstrates strong 5’➔3’ DNA polymerase activity with either DNA or RNA templates. We then performed quantitative polymerase chain reaction (qPCR) quantification for the three samples. We observed that cycle threshold (Ct) value of the Tn5 transposome sample is about 8 cycles smaller than the heat inactivated Tn5 sample or the control sample, indicating approximately 256 times more amplifiable products (see Figure 1f). We also tested different buffer conditions and found that the performance of Tn5 remained similar, indicating the robustness of the Tn5 tagmentation activity (see Figure S1c). Using Sanger sequencing, we validated that the adaptor sequences are indeed ligated to the insert sequences (see Figure S1d). Therefore, Tn5 can simultaneously fragment and ligate adaptors to both strands of RNA/DNA hybrids.

Having demonstrated the tagmentation activity of Tn5 on RNA/DNA hybrids, we then thought about its potential application. RNA/DNA duplexes can be found in many *in vivo* scenarios, including but not limited to R-loop and chromatin-bound lncRNAs (15, 16). Under *in vitro* conditions, RNA/DNA hybrids are also key intermediates in various molecular biology and genomics experiments. For instance, RNA has to be first reverse transcribed into cDNA in a traditional RNA-seq experiment so as to construct a library for sequencing. Because traditional RNA-seq library construction involves many laborious and time-consuming steps, including mRNA purification, fragmentation, reverse transcription, second-strand synthesis, end-repair and adaptor ligation, we attempted to replace the process using the tagmentation activity towards RNA/DNA duplexes. With the help of ATRAC-seq, these steps are replaced with a “one-tube” protocol (see Figure 2a), which uses total RNA as input material and involves just three seamless steps (reverse transcription, tagmentation and strand extension), without the need for a second strand synthesis step. We first conducted ATRAC-seq with 200 ng total RNA as input; we observed very high correlation in gene-expression levels among three replicates, indicating ATRAC-seq is highly reproducible (see Figure 2b). To test the robustness of ATRAC-seq, we performed the experiments with 20 ng and 2 ng total RNA. ATRAC-seq results are again highly reproducible among replicates (see Figure S2a, S2b). More importantly, gene expression level measured using different amount of starting materials remain consistent with each other (see Figure 2c).

**Figure 2.**
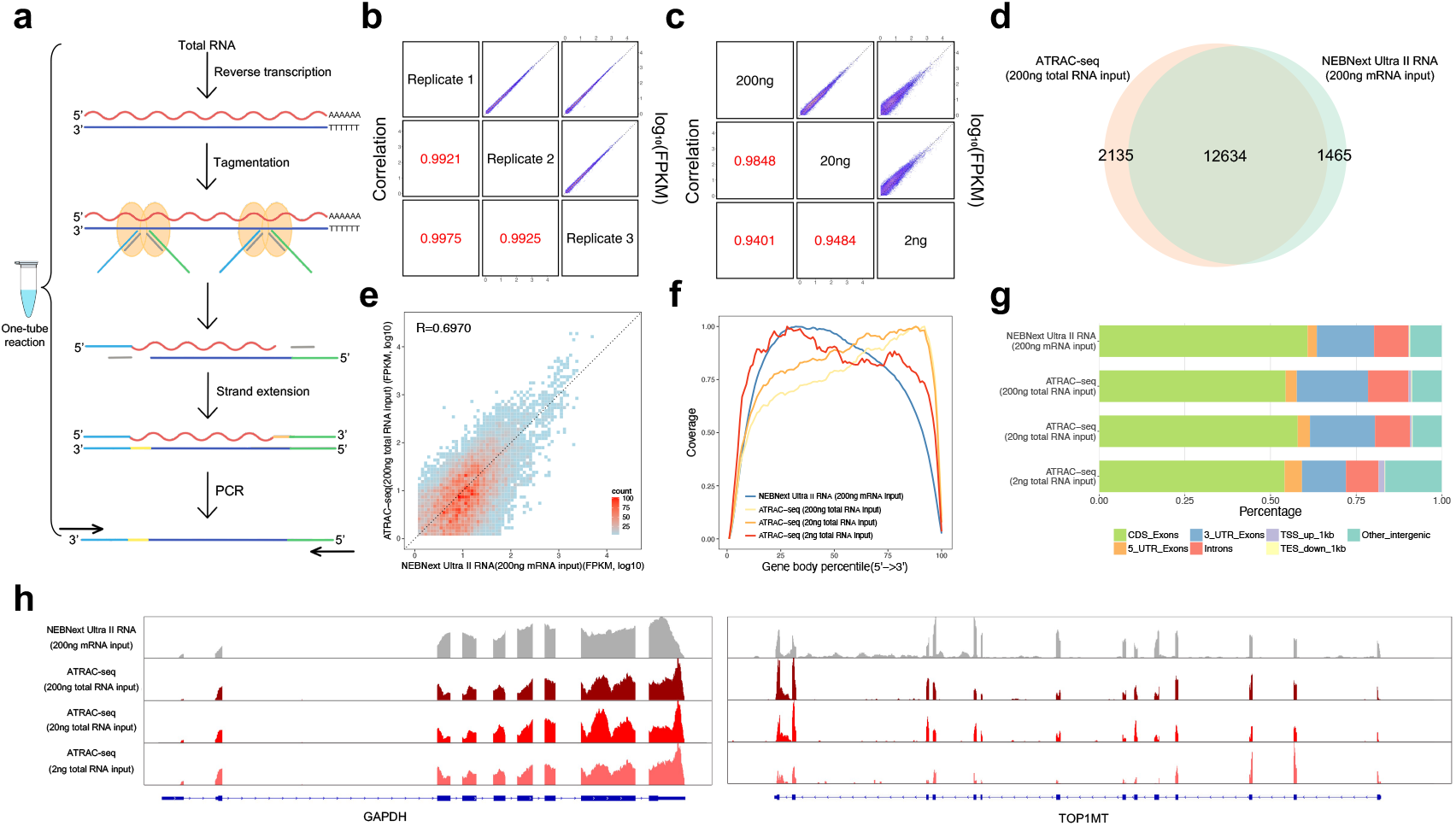
Workflow and evaluation of ATRAC-seq. **(a)** Workflow of ATRAC-seq. **(b)** Gene expression, measured by three technical replicates of ATRAC-seq with 200 ng total RNA as input, are shown as scatter plots in the upper right half. Pearson’s product-moment correlations are displayed in the lower left half. **(c)** Gene expression, measured by ATRAC-seq using 200 ng, 20 ng and 2 ng total RNA as input, are shown as scatter plots in the upper right half. Pearson’s product-moment correlations are displayed in the lower left half. **(d)** Venn Diagram of gene numbers detected by ATRAC-seq with 200 ng total RNA as input and NEBNext Ultra II RNA kit with 200 ng mRNA as input. **(e)** Scatterplot showing gene expression values for ATRAC-seq with 200 ng total RNA as input and NEBNext Ultra II RNA kit with 200 ng mRNA as input. Pearson’s product-moment correlation is displayed in the upper left corner. **(f)** Comparison of read coverage over gene body for ATRAC-seq with 200 ng, 20 ng and 2 ng total RNA as input and NEBNext Ultra II RNA kit with 200 ng mRNA as input. The read coverage over gene body is displayed along with gene body percentile from 5’ to 3’ end. **(g)** Comparison of the distribution of reads across known gene features for ATRAC-seq with 200 ng, 20 ng and 2 ng total RNA as input and NEBNext Ultra II RNA kit with 200 ng mRNA as input. **(h)** IGV tracks showing the coverage of two representative transcripts (GAPDH and TOP1MT). The data come from NEBNext Ultra II RNA kit and three sets of ATRAC-seq with different amount of total RNA.

We then compared the library quality between ATRAC-seq and NEBNext Ultra II RNA library prep kit, a commonly used kit for RNA-seq library construction. We found that ATRAC-seq libraries exhibited similar percentage of reads mapped to annotated transcripts, rRNA contamination and gene numbers to NEBNext data (see Table S1), despite the fact that ATRAC-seq directly uses total RNA as input material. Most of the genes detected by ATRAC-seq overlaps with that of NEBNext, with slightly more genes detected by ATRAC-seq (see Figure 2d). In addition, ATRAC-seq showed comparable performance to NEBNext in terms of gene expression measurement (see Figure 2e). Compared to NEBNext, the insert size of ATRAC-seq library was considerably shorter (see Figure S2c); nevertheless, we observed similar coverage distribution over gene body. ATRAC-seq also showed a slight tendency to 3’ end of the gene body (see Figure 2f). This 3’ bias of gene coverage decreased as the amount of starting materials reduced; hence it is likely due to incomplete reverse transcription of the 5’ end of transcripts when oligodT primers were used. Further inspection of reads distribution of ATRAC-seq over genome features revealed similar pattern for that of NEBNext (see Figure 2g). Coverages of some representative transcripts are shown in Figure 2h and Figure S2d.

To further investigate whether potential bias exists for ATRAC-seq, we compared the GC content of library prepared by ATRAC-seq with that of NEBNext. We found an enrichment of fragments with higher GC content in the ATRAC-seq libraries (see Figure S2e); whether or not this is due to the increased stability of GC-rich RNA/DNA hybrids, which is an asymmetric intermediate between A and B forms (17), remains to be demonstrated. Previous studies also found that Tn5 exhibits a slight insertion bias on dsDNA substrates (18–20). We thus characterized sites of Tn5-catalyzed adaptor insertion by calculating nucleotide composition of the first and last 10 bases of each sequencing read after adaptor trimming. Similar to dsDNA substrates, we also observed an apparent insertion signature on RNA/DNA hybrids (see Figure S2f). Nevertheless, per-position information contents were extremely low, suggesting such insertion bias is less likely to affect the uniformity of gene body coverage (see Figure S2g). Overall, when utilized as a library preparation method, ATRAC-seq demonstrates comparable performance with a traditional RNA library preparation method, but outcompetes the traditional method in terms of speed, convenience and cost.

## Discussion

Based on substrate diversity and the conserved catalytic domain of the retroviral integrase superfamily including the Tn5 transposase, we envision in this study that Tn5 may be able to directly tagment RNA/DNA hybrid duplexes, in addition to its canonical dsDNA substrates. Having validated such *in vitro* tagmentation activity, we developed ATRAC-seq, which enables one-tube, low-input and low-cost library construction for RNA-seq experiments. Compared to conventional RNA-seq methods, ATRAC-seq does not need to pre-extract mRNA and synthesize a second DNA chain after mRNA reverse transcription. Therefore, ATRAC-seq bypasses laborious and time-consuming processes, is compatible with low input, and reduces reagent cost. Collectively, these features enable library construction mediated by ATRAC-seq to competes the traditional methods.

Despite its unique advantages, there is room to further improve ATRAC-seq. For instance, ATRAC-seq exhibits signature at sites of adaptor insertion as well as a slight GC-bias for the insert sequences (see Figure S2e, S2f). Although we did not find a predominant motif and hence this signature does not appear to affect uniformity of coverage (see Figure S2g), it remains to be seen whether or not future engineered Tn5 mutants can bypass this bias. In fact, a Tn5 mutant showing reduced GC insertion bias on dsDNA has been reported previously (21). In addition, the *in vitro* tagmentation efficiency of Tn5 on RNA/DNA hybrids is low compared to its native substrate dsDNA. As wild-type Tn5 transposase has been engineered to obtain hyperactive forms (4, 22–24), it is also tempting to speculate that hyperactive mutants towards RNA/DNA hybrids could also be obtained through screening and protein engineering. Such hyperactive mutants are expected to have immediate utility in single-cell RNA-seq experiments, for instance. Moreover, Tn5 transposition *in vivo* has been harnessed to profile chromatin accessibility in ATAC-seq (10); it remains to be seen whether or not an equivalent version may exist to enable *in vivo* detection of R-loop, chromatin bound long non-coding RNA and epitranscriptome analysis (15, 16, 25). To summarize, ATRAC-seq manifests a “cryptic” activity of the Tn5 transposase as a powerful tool, which may have broad biomedical applications in the future.

## Materials and Methods

### Cell culture

HEK293T cells used in this study were daily maintained in DMEM medium (GIBCO) supplemented with 10% FBS (GIBCO) and 1% penicillin/streptomycin (GIBCO) at 37°C with 5% CO_2_.

### RNA isolation

Total RNA was extracted from cells with TRIzol (Invitrogen), according to the manufacturer’s instructions. The resulting total RNA was treated with DNase I (NEB) to avoid genomic DNA contamination. Phenol/chloroform extraction and ethanol precipitation were then performed to purify and concentrate total RNA. For mRNA isolation, two successive rounds of poly(A)+ selection were performed using oligo(dT)_25_ dynabeads (Invitrogen).

### Preparation of RNA/DNA hybrids

Total RNA, mRNA and an *in vitro* transcribed model mRNA (IRF9) were reverse transcribed into RNA/DNA hybrids by SuperScript IV reverse transcriptase (Invitrogen), according to the manufacturer’s protocol, with several modifications: 1) Instead of oligo d(T)_20_ primer, oligo d(T)_23_VN primer (NEB) was annealed to template RNA; 2) Instead of SS IV buffer, SS III buffer supplemented with 7.5% PEG8000 was added to the reaction mixture; 3) The reaction was incubated at 55°C for 2 h.

### Tn5 *in vitro* tagmentation on RNA/DNA hybrids

Partial double-stranded adaptor A and B were obtained by separately annealing 10 μM primer A (5’-TCGTCGGCAGCGTCAGATGTGTATAAGAGACAG-3’) and primer B (5’-GTCTCGTGGGCTCGGAGATGTGTATAAGAGACAG-3’) with equal amounts of mosaic-end oligonucleotides (5’-CTGTCTCTTATACACATCT-3’). Assembly of Tn5 with equimolar mixture of annealed Adaptor A and B was performed according to the manufacturer’s protocol (Vazyme). The resulting assembled Tn5 was stored at −20°C until use.

Tagmentation reaction was set up by adding RT products, 12 ng/μl assembled Tn5 and 1 U/μl SUPERase-In RNase Inhibitor (Invitrogen) to the reaction buffer containing 10 mM Tris-HCl (pH = 7.5), 5 mM MgCl_2_, 8% PEG8000. The reaction was performed at 55°C for 30 min, and then SDS was added to a final concentration of 0.04% and Tn5 was inactivated for 5 min at room temperature.

### Assays of tagmentation activity of Tn5 on RNA/DNA hybrids

For testing tagmentation activity of Tn5 on RNA/DNA hybrids, reactions were carried out as above, with 25ng mRNA derived RT products as substrate. The tagmentation products were then purified using 1.8X Agencourt RNAClean XP beads (Beckman Coulter) to remove Tn5 and excess free adaptors and eluted in 6μl nuclease-free water. The size distribution of RNA/DNA hybrids after tagmentation was assessed by a Fragment Analyzer Automated CE System with DNF-474 High Sensitivity NGS Fragment Analysis Kit (AATI).

For testing tagmentation activity of Tn5 on RNA/DNA hybrids by quantitative polymerase chain reaction (qPCR), tagmentation products purified as above (100X-diluted) was firstly strand-extended with 0.32 U/μl Bst 3.0 DNA Polymerase (NEB) and 1X AceQ Universal SYBR qPCR Master Mix (Vazyme) at 72°C for 15 min, and then Bst 3.0 Polymerase was inactivated at 95°C for 5 min. After adding 0.2 μM qPCR primers (5’-AATGATACGGCGACCACCGAGATCTACACTCGTCGGCAGCGTC-3’; 5’-CAAGCAGAAGACGGCATACGAGAT GTCTCGTGGGCTCGG-3’), qPCR was performed in a LightCycler (Roche) with a 5 min pre-incubation at 95°C followed by 40 cycles of 10 sec at 95°C and 40 sec at 60°C. For testing the effect of different buffers on tagmentation activity of Tn5 on RNA/DNA hybrids, buffers used were as follows: 1) Tagment buffer L (Vazyme); 2) Buffer with 8% PEG8000 (10 mM Tris-HCl at pH 7.5, 5 mM MgCl_2_, 8% PEG8000); 3) Buffer with 10% DMF (10 mM Tris-HCl at pH 7.5, 5 mM MgCl_2_, 10% DMF).

### ATRAC-seq library preparation and sequencing

For ATRAC-seq library preparation, all reactions were performed in one tube. Reverse transcription and tagmentation reactions were carried out as above. Strand extension reaction was performed by directly adding 0.32 U/μl Bst 3.0 DNA Polymerase and 1X NEBNext Q5 Hot Start HiFi PCR Master Mix (NEB) to tagmentation products and incubating at 72°C for 15 min, followed by Bst 3.0 DNA Polymerase inactivation at 80°C for 20 min. Next, 0.2 μM indexed primers were added to perform enrichment PCR as follows: 30 sec at 98°C, and then n cycles of 10 sec at 98°C, 75 sec at 65°C, followed by the last 10min extension at 65°C. The PCR cycles “n” depends on the amount of purified total RNA input (200 ng, n = 15; 20 ng, n = 20; 2 ng, n = 25). After enrichment, the library was purified twice using 1X Agencourt AMPure XP beads (Beckman Coulter) and eluted in 10 μl nuclease-free water. The concentration of resulting libraries was determined by Qubit 2.0 fluorometer with the Qubit dsDNA HS Assay kit (Invitrogen) and the size distribution of libraries was assessed by a Fragment Analyzer Automated CE System with DNF-474 High Sensitivity NGS Fragment Analysis Kit. Finally, libraries were sequenced on the Illumina Hiseq ×10 platform which generated 2 × 150 bp of paired-end raw reads.

### Data analysis

Raw reads from sequencing were firstly subjected to Trim_galore (v0.6.4_dev) (http://www.bioinformatics.babraham.ac.uk/projects/trim_galore/) for quality control and adaptor trimming. The minimal threshold of quality was 20, and the minimal length of reads to remain was set as 20 nt. Then trimmed reads were mapped to human (hg19) genome and transcriptome using Tophat2 (v2.1.1) (26), and the transcriptome was prepared based on the Refseq annotation of human (hg19) downloaded from the table browser of UCSC database. rRNA contamination were determined through directly mapping to the dataset of human rRNA sequence downloaded from NCBI by bowtie2 (v2.2.9) (27). Performances related to the processing of sam/bam file were done with the help of Samtools (v1.9) (28). The FPKM, gene body coverage, reads distribution, nucleotide composition for each position of read and GC content distribution of mapped reads were calculated by RseQC (v2.6.4) (29), and insert size of library was calculated by Picard Tools (v2.20.6) (http://broadinstitute.github.io/picard/). And all corresponding graphs were plotted using R scripts. Reads Coverage was visualized using the IGV genome browser (v2.4.16) (30).

## Acknowledgments

The authors would like to thank Mr. Dongsheng Bai for discussions. We thank National Center for Protein Sciences at Peking University in Beijing, China, for assistance with evaluation of tagmentation efficiency and library size distribution. This work was supported by the National Natural Science Foundation of China (nos. 31861143026 and 91740112 to C.Y.), and Ministry of Science and Technology of China (nos. 2019YFA0110900 and 2019YFA0802201 to C.Y.)

## Author Contributions

B.L., C.Z. and C.Y. conceived the project; B.L., L.D., D.Y. and C.Y. designed the experiments together and wrote the manuscript; B.L., L.D. and D.Y. performed experiments with the help of M.Z. and X.L.; B.L. performed the bioinformatics analysis with the help of C.Y. All authors commented on and approved the paper.

## Competing interests

The authors declare no competing interests.

## Data Availability

High-throughput sequence data has been deposited in Gene Expression Omnibus (GEO) under accession code GSE143422.

---

**Figure S1.**
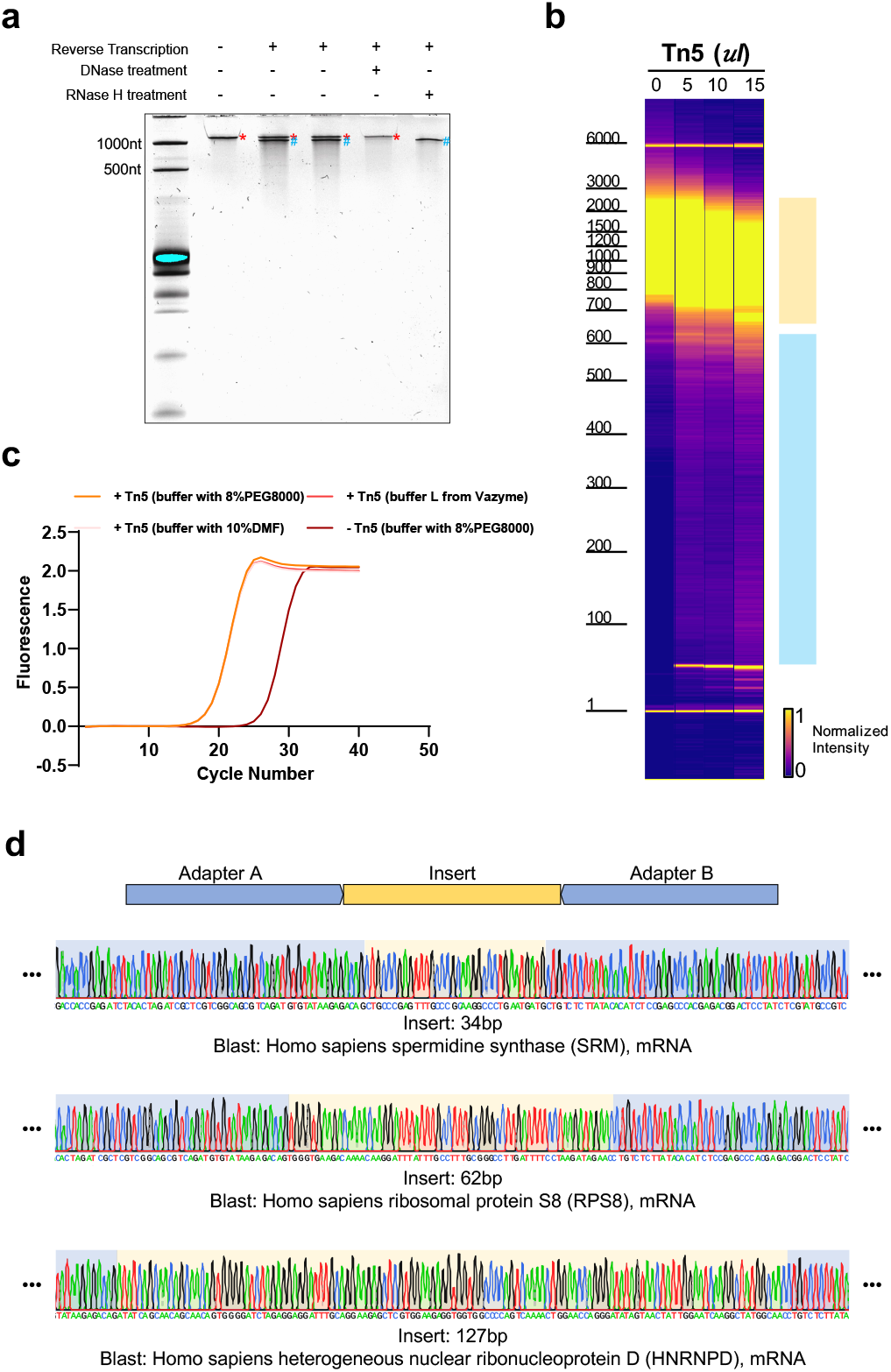
Tagmentation activity of Tn5 transposome on RNA/DNA hybrids. **(a)** Denaturing (8 M urea) polyacrylamide gel analysis of reverse transcription products of an *in vitro* transcribed mRNA (IRF9). Lane 1: ssRNA marker. Lane 2: *in vitro* transcribed mRNA (IRF9). Lane 3&4: reverse transcription products of an *in vitro* transcribed mRNA (IRF9). Lane 5: reverse transcription product treated with DNase I. Lane 6: reverse transcription product treated with RNase H. ssRNA and ssDNA is marked with a red asterisk and a blue pound sign, respectively. **(b)** Gel picture showing size distribution of RNA/DNA hybrids products of 50 μl reaction systems without Tn5 transposome, and with 5 μl, 10 μl, and 15 μl Tn5 transposome, respectively. The blue and orange patches denote small and large fragments, respectively. **(c)** qPCR amplification curve of tagmentation products without Tn5 treatment or with Tn5 treatment in three different buffers (see methods). Average Ct values are 26.41, 18.39, 18.33 and 18.34, respectively. **(d)** Sanger sequencing chromatograms of PCR products following RNA/DNA hybrid tagmentation and strand extension. Adaptor A and B sequences are highlighted with blue background color and insert sequences are highlighted with yellow background.

**Figure S2.**
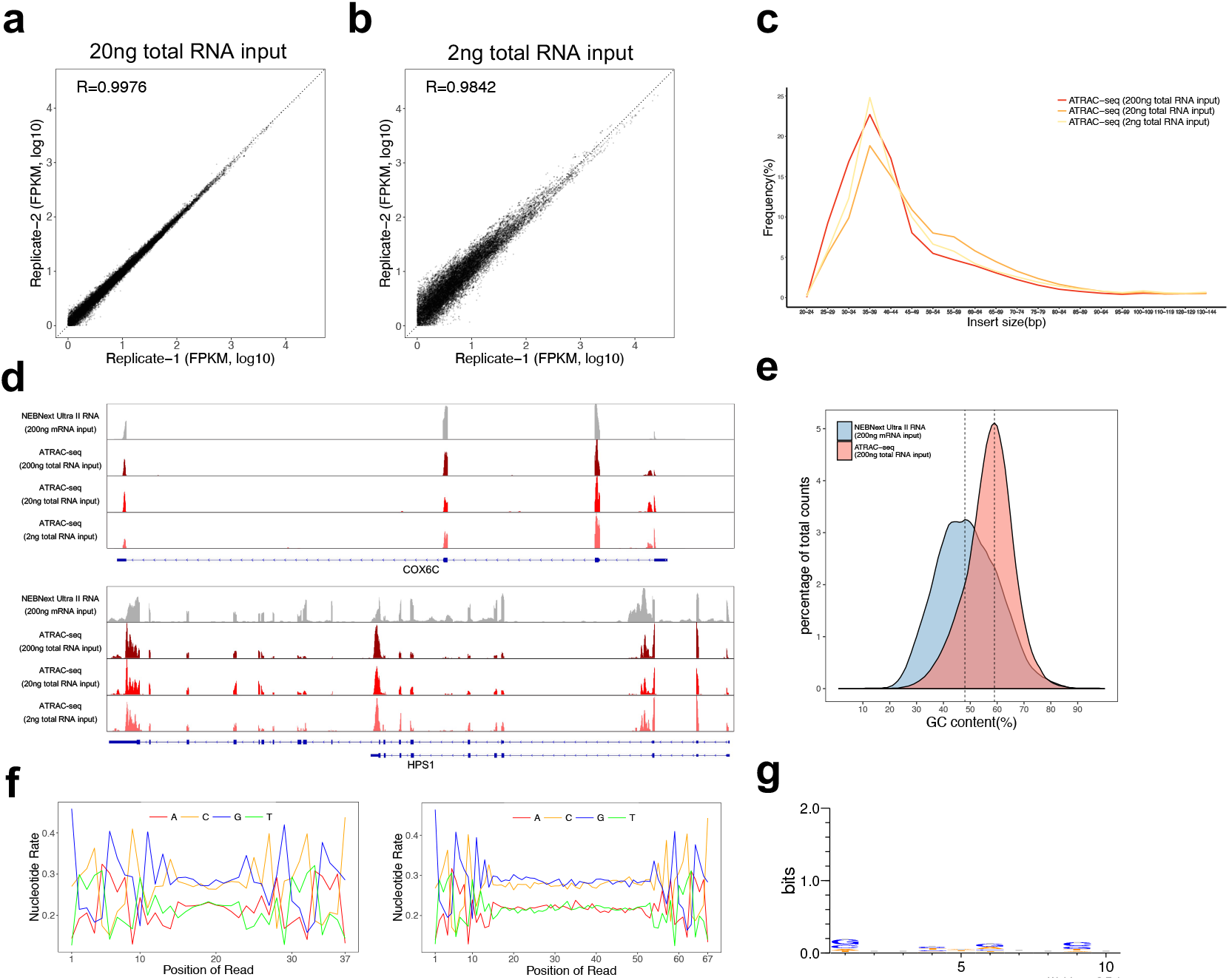
Quality assessment of ATRAC-seq. **(a)** Gene expression measured by two technical replicates of ATRAC-seq with 20 ng total RNA as input are shown as scatter plots. Pearson’s product-moment correlations are displayed in the upper left corner. **(b)** Gene expression measured by two technical replicates of ATRAC-seq with 2 ng total RNA as input are shown as scatter plots. Pearson’s product-moment correlations are displayed in the upper left corner. **(c)** Distribution of the insert size in ATRAC-seq data with 200 ng, 20 ng and 2 ng total RNA as input, respectively. **(d)** IGV tracks displaying the coverage of representative transcripts of a highly expressed gene COX6C, and a moderately expressed gene HPS1. **(e)** Distribution of GC content of all mapped reads from ATRAC-seq library with 200 ng total RNA as input and NEBNext Ultra II RNA library with 200 ng mRNA as input. Two vertical dashed lines indicate 48% and 59%. **(f)** Nucleotide versus cycle (NVC) plots showing percentage of observed bases at each position of mapped 37bp and 67bp reads from ATRAC-seq library with 200 ng total RNA as input. **(g)** Per-position information content of Tn5 insertion sites on RNA/DNA hybrids.

**Table S1.** Quality control of the sequencing results using NEBNext kit and ATRAC-seq.

| Library type | Mapping rate | rRNA rate | Genes |
| --- | --- | --- | --- |
| NEBNext Ultra II RNA<br>(200ng mRNA input) replicate 1 | 84.4% | 4.3% | 14099 |
| NEBNext Ultra II RNA<br>(200ng mRNA input) replicate 2 | 83.4% | 4.7% | 14072 |
| ATRAC-seq<br>(200ng total RNA input) replicate 1 | 87.9% | 9.8% | 14769 |
| ATRAC-seq<br>(200ng total RNA input) replicate 2 | 87.3% | 8.7% | 14713 |
| ATRAC-seq<br>(20ng total RNA input) replicate 1 | 89.3% | 5.5% | 14899 |
| ATRAC-seq<br>(20ng total RNA input) replicate 2 | 87.9% | 9.7% | 14964 |
| ATRAC-seq<br>(2ng total RNA input) replicate 1 | 69.5% | 20.6% | 13532 |
| ATRAC-seq<br>(2ng total RNA input) replicate 2 | 69.1% | 17.8% | 14213 |

